# ZNF146/OZF and ZNF507 target LINE-1 sequences

**DOI:** 10.1101/2021.05.24.445350

**Authors:** Kevin M. Creamer, Eric C. Larsen, Jeanne B. Lawrence

## Abstract

Repetitive sequences including transposable elements (TEs) and transposon-derived fragments account for nearly half of the human genome. While transposition-competent TEs must be repressed to maintain genomic stability, mutated and fragmented TEs comprising the bulk of repetitive sequences can also contribute to regulation of host gene expression and broader genome organization. Here we analyzed published ChIP-seq data sets to identify proteins broadly enriched on TEs in the human genome. We show two of the proteins identified, C2H2 zinc finger-containing proteins ZNF146 (also known as OZF) and ZNF507, are targeted to distinct sites within LINE-1 ORF2 at thousands of locations in the genome. ZNF146 binding sites are found at old and young LINE-1 elements. In contrast, ZNF507 preferentially binds at young LINE-1 sequences correlated to sequence changes in LINE-1 elements at ZNF507’s binding site. To gain further insight into ZNF146 and ZNF507 function, we disrupt their expression in HEK293 cells using CRISPR/Cas9 and perform RNA sequencing, finding modest gene expression changes in cells where ZNF507 has been disrupted. We further identify a physical interaction between ZNF507 and PRMT5, suggesting ZNF507 may target arginine methylation activity to LINE-1 sequences.

## Introduction

In the wake of whole genome sequencing, it has become clear that only a small portion of DNA codes for protein in higher organisms. While only about 1-2% of the roughly three billion bases in the human genome are protein-coding, an astounding 40-50% is derived from mobile repetitive sequences collectively known as transposable elements (TEs). Mobilization of TEs, sometimes referred to as “jumping genes,” poses a threat to host genome stability and is known to sometimes cause disease (Kazazian and Moran, 2017). In recent years, however, TEs and TE-derived sequences have been implicated as drivers of gene expression patterning, genome organization, development, and evolution in higher organisms (Hall et al., 2014; Lu et al., 2021; Rebollo et al., 2012).

Transposable elements multiply and spread in host genomes by “cut-and-paste” transposition of their DNA sequence or through an RNA intermediate (retrotransposition). Ultimately, to preserve genomic integrity TEs are largely inactivated by various mechanisms encoded by the host genome. Relatively few mobile TEs persist in the human genome, primarily consisting of a hundred or so long interspersed nuclear elements (LINE-1 or L1) and SVA (short interspersed nuclear element–VNTR–Alu) repeats that require L1 encoded reverse transcriptase activity to mobilize (Brouha et al., 2003; Wang et al., 2005). Instead, most TE DNA in the genome consists of degenerate remnants of prior mobilization events that have been truncated or mutated and no longer harbor the activity necessary to “jump.”

Transposition-competent TEs are generally inactivated at the level of transcription. TEs sometimes, however, escape this repression. In humans, L1s mobilize in the germline and are a major source of inheritable structural variation between individuals (Ewing and Kazazian, 2010; Mir et al., 2015). Recent studies have revealed that L1 elements are also activated in specific contexts outside of the germline, such as stress, cancer, and neural development, with disease-amplifying or functional implications (Burns, 2017; Coufal et al., 2009; Faulkner and Garcia-Perez, 2017; Kazazian and Moran, 2017).

Irrespective of their mobility, abundant TE sequences are non-randomly distributed in host genomes and contribute to development and gene expression patterning through recognition by DNA-binding proteins (Bourque et al., 2008; Korenberg and Rykowski, 1988; Manuelidis and Ward, 1984; Rebollo et al., 2012). For instance, HERV-H/MERV-L TEs are known to be expressed early during embryogenesis and are important regulators of pluripotency and embryogenesis by regulating the expression of neighboring genes, a process driven by sequence-specific recognition of TE DNA by transcription factors (TFs) (Kigami et al., 2003; Robbez-Masson and Rowe, 2015; Wang et al., 2014). L1 elements are also known to be highly expressed in the early mouse embryo and contribute to proper embryonic development (Fadloun et al., 2013; Jachowicz et al., 2017).

While several DNA-binding proteins are known to target TEs, the complex picture of TF binding to TE sequences remains incomplete and the functional significance of these interactions on cell or organism biology is understudied, and largely unknown. Here we analyze published ChIP-seq data sets to identify proteins that broadly recognize TE-derived sequences in the human genome. We focus on two poorly studied zinc finger proteins, ZNF146 (also known as OZF) and ZNF507, and characterize their recognition motifs within L1 repeats. Interestingly, we find ZNF146 is a highly conserved protein that retains targeting to thousands of relatively old L1 sequences in the genome. In contrast, ZNF507 was only observed at relatively young L1 sequences. We go on to functionally test the impact depletion of these proteins has on the transcriptome of HEK293 cells. Finally, we identify an interaction between ZNF507 and PRMT5 by co-immunoprecipitation and mass spectrometry, yielding potential insight into the function of ZNF507 in the cell.

## Results

### Identification of DNA-binding proteins that target TEs

In order to identify proteins that broadly target transposable elements (TEs) we developed a strategy to test chromatin immunoprecipitation with sequencing (ChIP-seq) data sets for enrichment on TEs using a simple peak intersection strategy (Figure 1A). For this study we analyzed all ENCODE transcription factor (TF) ChIP-seq experiments (as of March 2020) for which peaks had been identified, spanning a large number of transcription factors, cell types, and sequencing methodologies (Consortium, 2012; Imbeault et al., 2017). Peaks were then intersected with Repeatmasker annotated repeats for the most abundant TE classes in the human genome. Frequency of intersection between peaks for each experiment was then compared to frequency of intersection with background models generated by random shuffling to generate a ratio of observed elements relative to what would be expected by random chance (Supplemental Table 1, Figure 1B).

**Figure 1.**
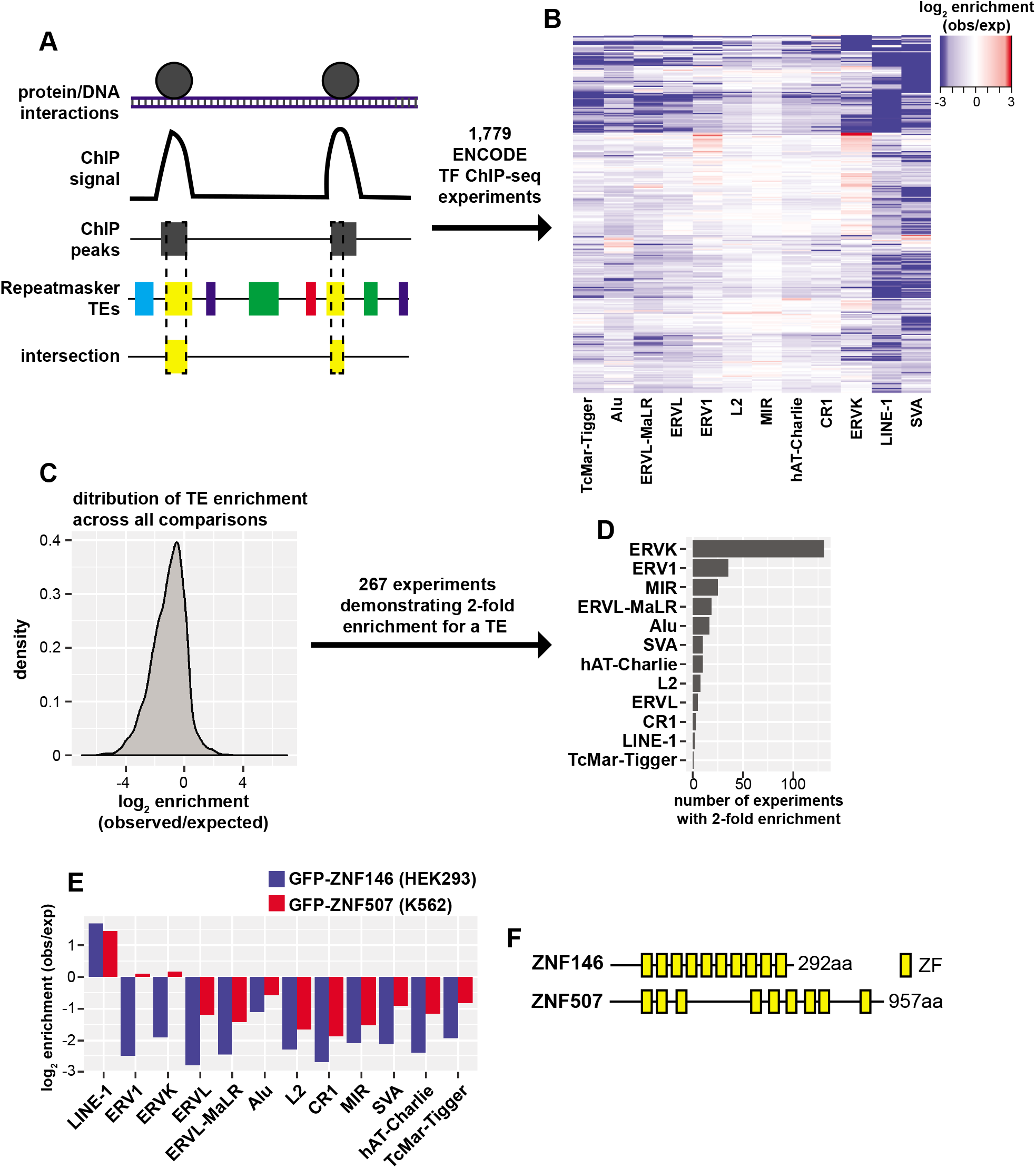
Identification of TFs that transposable elements. (A) Schematic of ENCODE TF ChIP-seq peak intersection with Repeatmasker annotated TEs. (B) Heatmap of TF ChIP-seq enrichment on abundant TE classes calculated as the log2 ratio of observed (obs) peak/TE intersections relative to simulated shuffled peaks (expected, exp). (C) Density of enrichment across all ChIP-seq experiment peak and TE class comparisons. (D) Bar chart of the number of TF ChIP-seq experiments demonstrating 2-fold enrichment binned by TE class. (E) Bar chart of peak enrichment on TE classes for GFP-ZNF146 ChIP-seq in HEK293 cells and GFP-ZNF507 in K562 cells. (F) Schematic representation of ZNF146 and ZNF507 proteins and the distribution of their C2H2 zinc finger domains.

While most peak/TE comparisons indicated TF binding to TEs occurred less frequently than expected by random chance (Figure 1C), 267 experiments indicated peak enrichment of greater than 2-fold (observed/expected) for at least one TE class (Figure 1D). Interestingly, enrichment was most frequently (>100 experiments) observed for ERVK transposons. We were additionally struck with the lack of TFs identified by this approach targeting LINE-1 elements. Just two experiments, ChIP-seq for GFP-ZNF146 in HEK293 and GFP-ZNF507 in A529 cells demonstrated 2-fold enrichment for LINE-1 elements. This enrichment was specific for LINE-1 as these proteins were not associated with other TE classes (Figure 1E).

ZNF146 (also known as OZF or Only Zinc Fingers) and ZNF507 genes are both found on chromosome 19 and encode proteins with 10 and 9 C2H2 zinc finger motifs, respectively; neither protein has any other easily identifiable domains (Figure 1F). Zinc finger motifs can contribute to protein-protein and protein-RNA interactions, but are best studied for their contribution to transcription factor binding to specific DNA sequences (Laity et al., 2001). ZNF146/OZF was previously identified as being overexpressed in certain cancers (Ferbus et al., 1999; Ferbus et al., 2003). Still, neither ZNF146 or ZNF507 have been extensively studied and little is known about their function despite being predicted to be widely expressed in human cells (Uhlen et al., 2015; Uhlen et al., 2017).

LINE-1 sequences are widely distributed and highly abundant in the human genome. Most L1 sequences are truncated and degenerate, with varying divergence from transposon-competent consensus sequences. There are estimated to be just 100 transposition-competent L1 elements, 4,000 mutated but full-length elements, and several hundred thousand truncated L1 fragments in the human genome (Brouha et al., 2003). Because of LINE-1 prevalence in the human genome, the small number and specificity of TFs identified by this approach, and the understudied nature of the two proteins identified, we decided to characterize the targeting of ZNF146 and ZNF507 to L1 elements in greater detail.

### ZNF146 and ZNF507 target LINE-1 ORF2

The above analyses indicated ZNF146 and ZNF507 preferentially bind to LINE-1 sequences. In order to verify these observations, we performed additional in-depth analysis of ENCODE’s ChIP-seq experiments. Consistent with L1 targeting, peaks for both ZNF146 and ZNF507 ChIP-seq experiments were found mostly (81% and 76% of peaks, respectively) within annotated L1 elements (Figure 2A). Despite high frequency of targeting to L1 elements for both proteins, ZNF146 peaks and ZNF507 peaks themselves did not frequently coincide (Figure 2B), suggesting they target different sequences within L1 elements.

**Figure 2.**
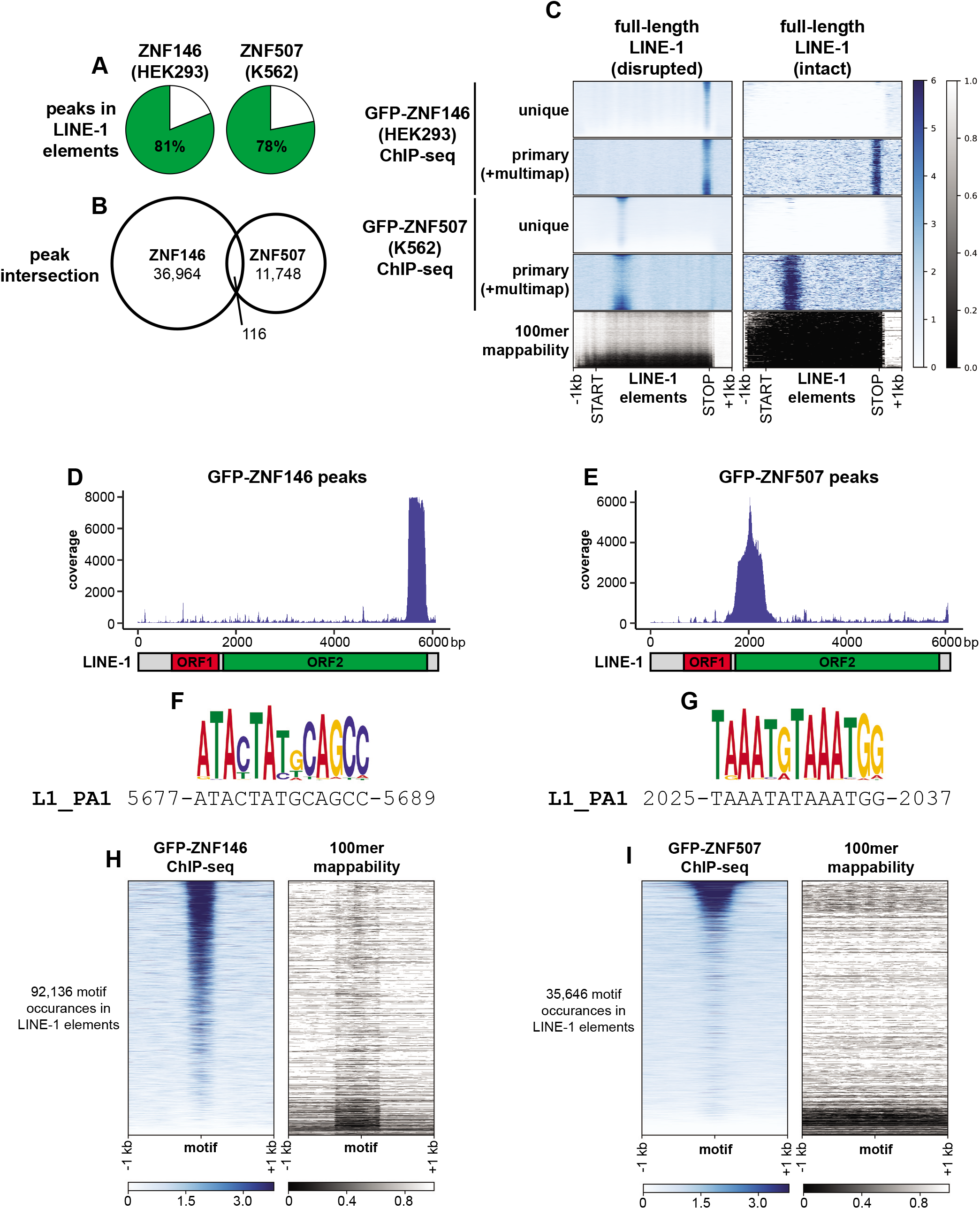
ZNF146 and ZNF507 target LINE-1 sequences. (A) Pie chart representing the percentage of peaks in the indicated ChIP-seq experiments overlapping annotated LINE-1 elements. (B) Ven diagram of peak interval intersections between the indicated ChIP-seq experiments. (C) Meta-analysis of ChIP-seq enrichment at full-length disrupted (n=13,092) or intact (ORF2 or ORF1 and ORF2) LINE-1 elements (n=245) normalized to 5kb. Below is the 100mer mappability score for the same regions (lower scores indicate lower mappability) (D,E) Coverage of L1-mapping reads directly mapped to LINE-1 (human specific, PA1) consensus sequence. Below is a schematic of full-length LINE-1. (F,G) Enriched motifs (see methods) and corresponding LINE-1 (human specific, PA1) consensus sequence determined from ZNF146 and ZNF507 ChIP-seq experiments, respectively. (H,I) Heatmap of ChIP-seq enrichment centered around occurrences of identified motifs (FIMO, p < 5e-6) found in annotated LINE-1 elements in their genomic context. On the right is the theoretical mappability of 100bp reads in the same regions.

To test this idea we performed meta-analysis of ChIP-seq coverage for both proteins at fulllength L1 elements (Figure 2C). Due to their repetitive nature, full-length L1 elements are often difficult to analyze by sequencing methods, however for these experiments ENCODE performed 100bp paired-end sequencing, allowing for relatively high mappability at disrupted (degenerate or mutated), full-length elements (Figure 2C, left panels). In contrast, intact (retaining coding potential) L1s had very low mappability (Figure 2C, right panels).

Using uniquely mapping reads, concentrated enrichment of ChIP-seq signal was observed for both ZNF146 and ZNF507 at disrupted, full-length L1s, with ZNF507 enrichment most prominent toward the 5’ end and ZNF146 toward the 3’ end of these elements. Using primary reads (which allow for multimapping reads to be randomly assigned to one location), similar enrichments could be observed on intact L1 elements. Hence while it is not possible to ascertain whether ZNF146 and ZNF507 bind at individual, intact elements, these proteins likely target intact L1s as well.

We then resolved ZNF146 and ZNF507 binding by calculating coverage after performing local alignment of peak sequences to the L1PA1 (human specific) consensus sequence. LINE-1 transposons code for two proteins, ORF1p and ORF2p, both of which are required for transposition of LINE-1 (Moran et al., 1996). ORF1 encodes a chaperone while ORF2 encodes a reverse transcriptase essential for transposition of not only L1, but other non-autonomous transposable elements as well (Dewannieux et al., 2003; Kolosha and Martin, 1997; Mathias et al., 1991; Ostertag et al., 2003). ZNF146 and ZNF507 peaks specifically aligned to the 3’ and 5’ ends of the ORF2 coding region of L1, respectively, with no enrichment within ORF1 (Figures 2D and 2E).

We further identified likely binding motifs within L1 ORF2 by performing motif discovery (Figures 2F and 2G). The most significant motif identified for each ChIP-seq experiment closely matched sequences in the L1PA1 consensus sequence at positions of highest ChIP-seq enrichment (Figures 2D-G). We then located all instances of the determined motifs for ZNF146 and ZNF507 in L1 elements genome-wide and performed meta-analysis of ChIP-seq signal at these motifs. As expected, robust ChIP-seq signal was found centered around these motifs for both proteins (Figures 2H and 2I). Thus, targeting of ZNF146 and ZNF507 to L1 elements appears to be genuine and occurs at opposite ends of the ORF2 coding region.

### ZNF507 preferentially binds at young LINE-1 sequences

While ChIP-seq signal enrichment was observed for ZNF146 at nearly all mappable instances of its predicted binding sites (Figure 2H), ZNF507 ChIP-seq signal was enriched at a fraction of its motif sites, suggesting that ZNF507 may only bind to a subset of L1 elements. LINE-1 sequences in the genome have been classified based on diagnostic sequence variants. Comparative genomics and expected sequence divergence over time have further allowed for approximate ageing of L1 subfamilies containing these shared sequence variants from youngest (L1PA1, human specific) to oldest (L1PA17) primate-specific LINE-1 elements (Giordano et al., 2007; Khan et al., 2006; Konkel et al., 2010; Smit et al., 1995).

To determine whether ZNF146 or ZNF507 might differentially recognize L1 subfamilies we binned motif instances in LINE-1 elements by subfamily, and performed meta-analysis of ChIP-seq signals around these motifs. As expected from unbinned meta-analysis (Figure 2H), ZNF146 ChIP-seq enrichment was observed at nearly all predicted mappable binding sites in all subfamilies (Figure 3A, upper panels). Because the youngest subfamilies (L1PA1 and L1PA2) are less uniquely mappable, binding to those families could not be reliably determined. We note, however, that those families contain the same conserved binding motif and enrichment can be observed at young, intact elements when including multimapping reads (Figure 2C), suggesting ZNF146 likely binds at these sequences as well.

**Figure 3.**
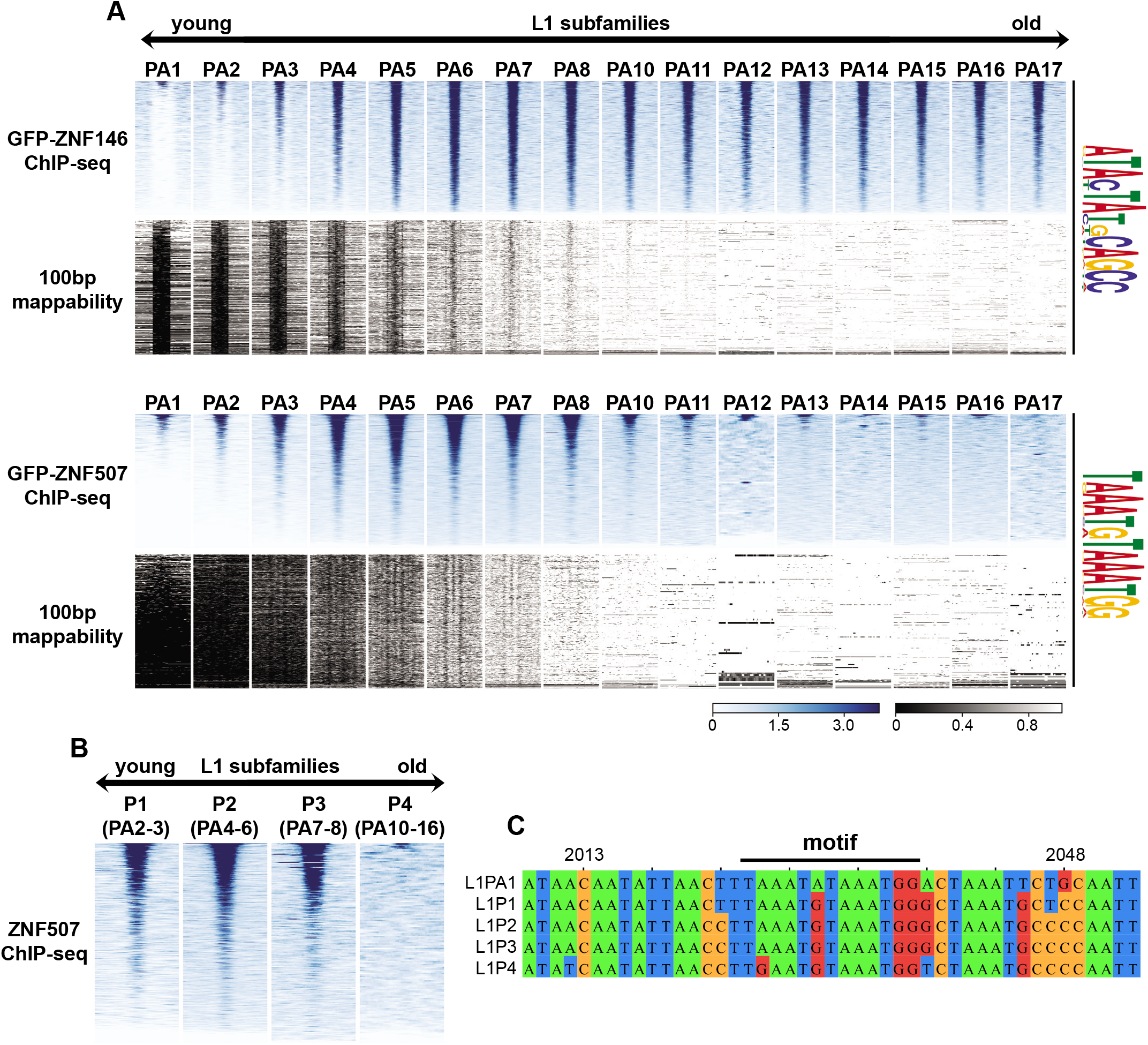
ZNF507 does not bind at old L1PA elements. (A) Heatmap of ChIP-seq enrichment centered around occurrences of identified motifs (FIMO, p<5e-6) +/-1Kb after intersection and binning by LINE-1 subfamily. Below the ChIP-seq heatmaps (blue) is the theoretical mappability of 100bp reads for the same regions (greyscale). (B) Heatmaps as above but for LINE-1 subfamilies binned by their ORF2-derived classification. Relationship between the ORF2 and 3’UTR classifications is given in parentheses. (C) Multiple sequence alignment of L1 subfamily consensus sequences. Positions are derived from the L1PA1 consensus sequence. Position of the ZNF507 ChIP seq binding motif identified by MEME is indicated above the alignment.

In contrast, we observed strong enrichment of ZNF507 only in young subfamilies, particularly in L1PA3-L1PA8 (Figure 3A, lower panels). Again, while binding at L1PA1 and L1PA2 sequences could not be definitively determined due to mappability, binding of ZNF507 was observed at intact L1 sequences when allowing for multimapping (Figure 2C), suggesting ZNF507 likely binds at these elements as well. Enrichment was essentially absent or rarely observed for ZNF507 in older subfamilies, particularly PA12-PA17. L1 classifications are determined largely by 3’ UTR sequence variations (as above) but are sometimes classified using upstream ORF2 sequences (L1P1, youngest, to L1P4, oldest). ZNF507 enrichment was clearly observed at motifs in L1P1, L1P2, and L1P3 subfamilies, but essentially absent at motifs in L1P4 elements (Figure 3B). Interestingly the L1P4 consensus sequence deviates from younger subfamilies within the core motif we identified as well as two other nearby residues, perhaps explaining the lack of binding observed for ZNF507 at these elements (Figure 3C).

Zinc finger proteins and TEs are in some cases thought to have co-evolved in an “evolutionary arms race” of natural selection (Cosby et al., 2019; Jacobs et al., 2014). Since ZNF507 binding was observed only at young L1 subfamilies, we wondered if this specificity change could be related to amino acid substitutions in ZNF507 zinc fingers. Interestingly, multiple sequence alignment of ZNF507 ZFs revealed three separate residues in ZNF507 ZFs that are highly conserved among higher primates (apes and monkeys) but were not found in lower primates and other mammals (Figure 4A). The timing of these substitutions appeared to closely parallel changes in L1 sequence and differences in ZNF507 binding (compare timeline in Figures 4B to Figures 3A, 3B, and 3C), collectively suggesting a relationship between the amino acid substitutions in ZNF507 ZFs and L1 targeting specificity.

**Figure 4.**
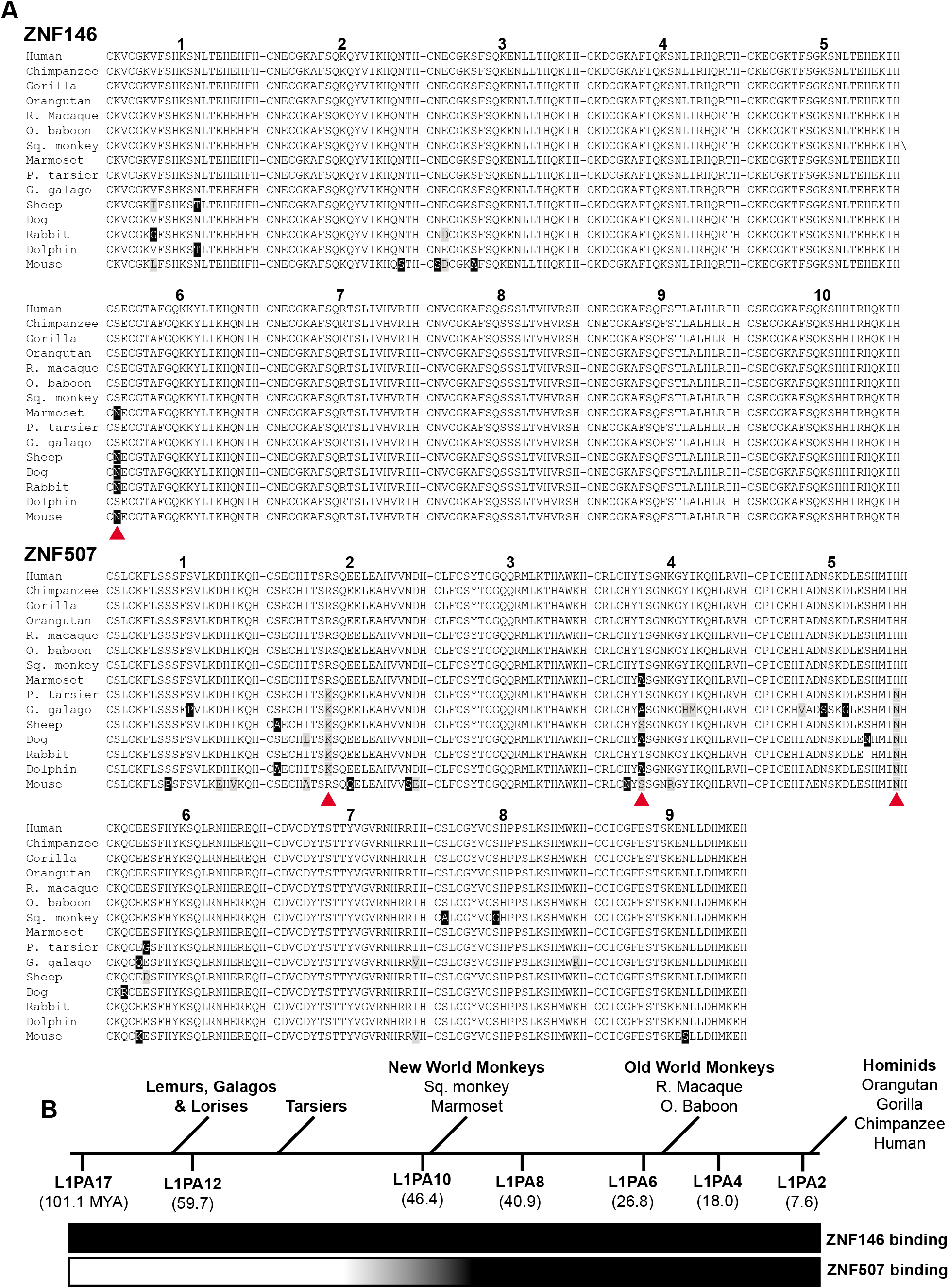
Conservation of ZNF146 and ZNF507 zinc finger domains. (A) Alignment of protein sequences for ZNF146 and ZNF507 zinc finder domains (indicated by number above alignment) in candidate mammals. Abbreviated common names are used (R. macaque, Rhesus macaque; O. baboon, Olive baboon; Sq. monkey, Squirrel monkey; P. tarsier, Philippine tarsier; G. galago, Garnett’s galago). Black boxes indicate residue differences from human proteins. Grey boxes indicate nonidentical but similar residues. Red arrows indicate residues conserved in monkeys and apes, but not frequently observed in tarsiers, lower primates, or other mammals. (B) Approximate timeline of primate evolution relative to the age of L1PA subfamilies (millions of years ago, MYA), adapted from Khan et al. and Konkel et al. Qualitative assessment of ZNF146/ZNF507 binding to L1 subfamilies (inferred from ChIP-seq) is displayed as a gradient below from non-binding (white) to binding (dark).

### Impact of ZNF146 and ZNF507 depletion on a somatic cell transcriptome

The above results demonstrate ZNF146 and ZNF507 target thousands of L1 sequences in the genome. Binding at these regions could potentially influence L1 or endogenous gene expression. We therefore disrupted ZNF146 and ZNF507 expression in human embryonic kidney cells (HEK293) using CRISPR/Cas9 to determine what impact depleting these proteins can have on the transcriptome of a somatic cell line.

Cells were transiently transfected with Cas9 and sgRNA expressing plasmids targeting either gene or a non-empty vector control (Ran et al., 2013). After transient selection and outgrowth, whole cell extracts were analyzed by western blot to confirm target gene disruption. Figure 5A shows ZNF146 and ZNF507 were essentially undetectable in Cas9-targeted cells (Figure 5A), which had no obvious growth or morphology defects. We then isolated RNA and performed transcriptomic analysis by paired-end RNA sequencing following ribosome depletion.

**Figure 5.**
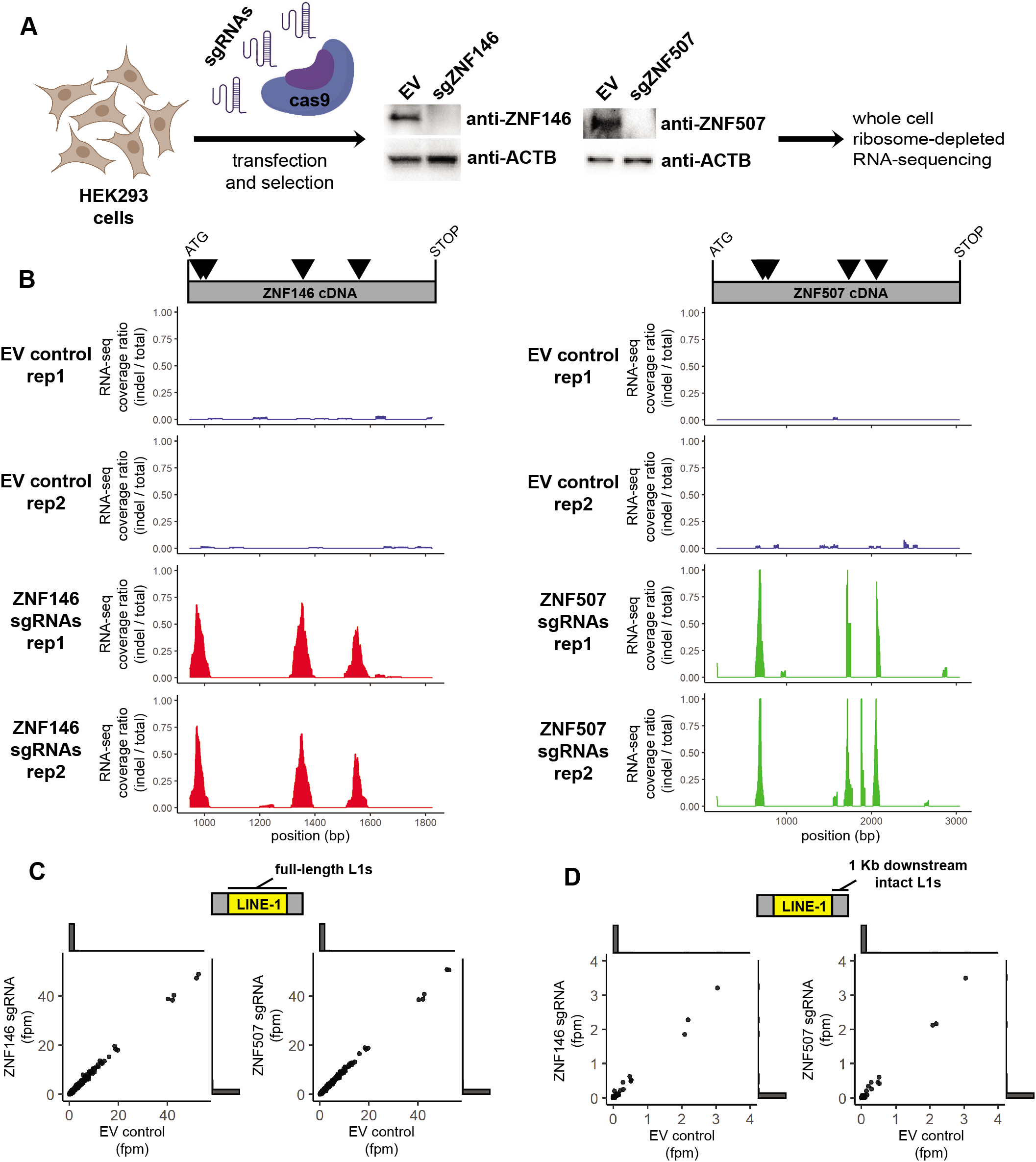
ZNF146 and ZNF507 are not required to silence LINE-1 elements in HEK293 cells. (A) Schematic of CRISPR/cas9 disruption approach. HEK293 cells were transiently transfected with spCas9-sgRNA-puro plasmids with either sgRNAs targeting ZNF146 or ZNF507 respectively or an empty vector (EV) control. Cells were briefly selected with puromycin before outgrowth. Fourteen days post-transfection cells were harvested for western blot analysis (middle panels) to assess disruption of protein expression. RNA was isolated for RNA-sequencing analysis. Disruption and RNA-sequencing were performed in duplicate. (B) RNA-sequencing coverage was calculated for reads mapping to ZNF146 (left) or ZNF507 cDNA (right) after filtering for reads marked with or without indels during alignment. Data is presented as the ratio of coverage calculated for reads with indels (determined by CIGAR tags) over total reads in each RNA-sequencing replicate. Triangles in the schematic above represent sgRNA targets. (C) Comparison of RNA-sequencing depth in fragments per million (fpm) mapping to individual full-length LINE-1 elements. Outside of the scatter plot are histograms showing most values are at or near zero. (D) Comparison of RNA-sequencing depth in fragments per million (fpm) mapping to 1 Kb regions downstream of intact (ORF1/ORF2 or ORF2 only) LINE-1 elements. Outside of the scatter plot are histograms indicating most values are at or near zero. Schematics created with BioRender.com.

First, we confirmed Cas9 targeting by aligning RNA-seq reads to ZNF146 and ZNF507 cDNA reference sequences using Tophat2 and BWA aligners to rescue unmapped reads as previously described (Deininger et al., 2017). We then compared the coverage ratio of reads marked as having insertions or deletions (indels) during alignment versus reads without indels. For both ZNF146 and ZNF507 indels were frequently detected at sgRNA target sites that were not observed in control cells, consistent with efficient disruption and protein depletion observed by western blot (Figure 5B).

We then assessed whether we could detect an increase in RNA originating from full-length L1 elements. Full-length and intact L1 elements are known to be heavily repressed in most cell lines. Because of this and their repetitive nature, the vast majority of full-length elements were not detected using uniquely mapping RNA-seq reads (see histogram of counts at or near zero in Figures 5C). For those that were detectable we did not observe any that were differentially expressed (p < 0.05) in cell populations in which ZNF146 or ZNF507 were disrupted (Figure 5C).

Although intact LINE-1 elements are themselves highly unmappable by unique reads, their expression can reliably be detected in RNA-seq experiments by quantifying reads mapping to immediately downstream regions which become expressed as a result of read-through transcription (Philippe et al., 2016). Similar to what we observed within L1 elements, few RNA-seq reads mapped downstream of intact L1 elements, consistent with their general repression (Figure 5D). For those downstream regions for which we did observe low-level expression, transcripts from these regions were not differentially expressed when ZNF146 or ZNF507 were disrupted. Although we can not rule out increased levels of rare or stochastic L1 activation, these results indicate L1 elements are not broadly activated in response to ZNF146 and ZNF507 depletion in HEK293 cells.

ZNF146 and ZNF507 ChIP-seq peaks are not restricted to intergenic regions as we also observed many peaks for both proteins in regulatory and genic regions including at annotated enhancers and promoters (Figure 6A). Additionally, a large numbers of peaks (>50%) for both ZNF146 and ZNF507 were located in introns. Targeting to these regulatory and genic regions could potentially influence endogenous gene expression.

**Figure 6.**
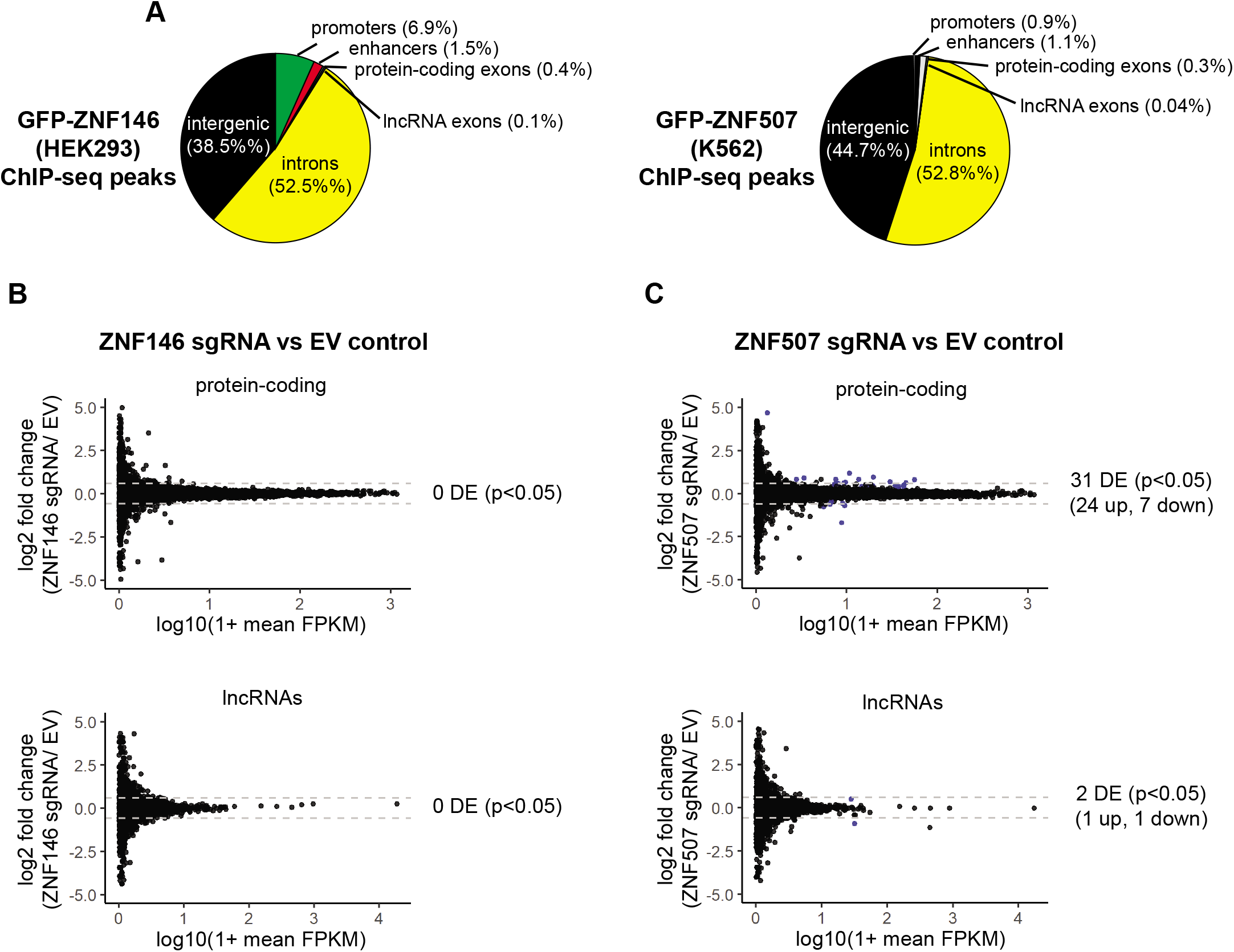
Disruption of ZNF507 leads to modest changes in gene expression. (A) Pie chart of ChIP-seq peak intersection with genomic features. Promoters and enhancers are PHAMTOM-annotated. Genic features are RefSeq-annotated. (B,C) Comparison of RNA-sequencing depth in fragments per kilobase per million mapped reads (FPKM) mapping to protein-coding transcripts (top) or lncRNAs (bottom). ZNF146 (left) or ZNF507 (right) disrupted cells are compared to EV control. Dotted grey lines mark 1.5-fold change. Transcripts which were differentially expressed (p < 0.05) are colored blue.

We therefore assessed what impact, if any, loss of ZNF146 or ZNF507 had on gene expression. Remarkably, despite observed binding peaks at over 40,000 sites in ChIP-seq data, we did not observe differential expression of any protein-coding or lncRNA (p < 0.05) in cells depleted of ZNF146 in HEK293 cells (Figure 6B). Interestingly, however, ZNF507 disruption resulted in the differential expression of 30 protein-coding (24 up and 6 down-regulated) and two lncRNA (one up and one down-regulated) transcripts, indicating depletion of ZNF507 influenced non-retroviral gene expression (Figure 6C, Table 1).

**Table 1.** Differentially expressed protein-coding genes after ZNF507 disruption.

| ENSEMBL ID | symbol | log2 fold change<br>(sgZNF507/EV) | padj* | chromosome |
| --- | --- | --- | --- | --- |
| ENSG00000198715 | GLMP | -1.705643082 | 6.99E-29 | chr1 |
| ENSG00000103174 | NAGPA | 1.169592472 | 4.40E-22 | chr16 |
| ENSG00000110442 | COMMD9 | 0.947934187 | 5.24E-18 | chr11 |
| ENSG00000122965 | RBM19 | 0.689678085 | 2.15E-16 | chr12 |
| ENSG00000059691 | GATB | 0.821869765 | 3.65E-16 | chr4 |
| ENSG00000102393 | GLA | 0.81066298 | 2.52E-15 | chrX |
| ENSG00000185896 | LAMP1 | 0.637840578 | 7.58E-15 | chr13 |
| ENSG00000090581 | GNPTG | 0.818300413 | 6.83E-13 | chr16 |
| ENSG00000167716 | WDR81 | -0.690739581 | 1.18E-10 | chr17 |
| ENSG00000124839 | RAB17 | 4.677234569 | 3.18E-10 | chr2 |
| ENSG00000160695 | VPS11 | 0.6614881 | 1.20E-09 | chr11 |
| ENSG00000168813 | ZNF507 | -0.613938125 | 2.11E-09 | chr19 |
| ENSG00000198908 | BHLHB9 | -0.758590795 | 3.57E-08 | chrX |
| ENSG00000103042 | SLC38A7 | 0.611608645 | 5.06E-08 | chr16 |
| ENSG00000176994 | SMCR8 | 0.457690189 | 2.37E-07 | chr17 |
| ENSG00000197081 | IGF2R | 0.416262343 | 2.75E-07 | chr6 |
| ENSG00000103249 | CLCN7 | 0.457423373 | 1.32E-06 | chr16 |
| ENSG00000157593 | SLC35B2 | 0.494377197 | 1.43E-06 | chr6 |
| ENSG00000106266 | SNX8 | -0.545579961 | 1.82E-06 | chr7 |
| ENSG00000047249 | ATP6V1H | 0.570565355 | 2.76E-06 | chr8 |
| ENSG00000064601 | CTSA | 0.487320189 | 3.47E-06 | chr20 |
| ENSG00000144455 | SUMF1 | 0.836597857 | 5.32E-06 | chr3 |
| ENSG00000159720 | ATP6V0D1 | 0.394188331 | 5.88E-06 | chr16 |
| ENSG00000244045 | TMEM199 | 0.590532873 | 6.49E-06 | chr17 |
| ENSG00000087088 | BAX | 0.414954399 | 8.94E-06 | chr19 |
| ENSG00000242802 | AP5Z1 | 0.525924987 | 9.58E-06 | chr7 |
| ENSG00000198356 | ASNA1 | 0.447441419 | 1.71E-05 | chr19 |
| ENSG00000169682 | SPNS1 | 0.896876564 | 2.05E-05 | chr16 |
| ENSG00000213614 | HEXA | -0.448754497 | 2.14E-05 | chr15 |
| ENSG00000103043 | VAC14 | 0.401200129 | 2.46E-05 | chr16 |

\*Differentially expressed genes calculated by DESeq2. Cutoff $p < 0.05$ .

### Identification of a physical interaction between ZNF507 and PRMT5

Transcription factors frequently regulate gene expression by recruiting chromatin modifying complexes to regulatory sequences. Hence, we wondered whether ZNF507 might associate with a known chromatin modifying enzyme. To test this idea, we expressed 3xFLAG-tagged ZNF507 in HEK293 cells by transient transfection (Figure 7A) and performed coimmunoprecipitation followed by unbiased interaction discovery by liquid chromatography with tandem mass spectrometry (LC-MS/MS). FLAG-tagged ZNF507 localized as expected to the nucleus in HEK293 cells (Figure 7B). Immunoprecipitation (IP) was performed on cells expressing 3xFLAG-ZNF507 as well as an empty vector control using anti-FLAG beads. Proteins were eluted with 3xFLAG peptide and partially resolved by SDS-PAGE prior to proteomic analysis by LC-MS/MS.

**Figure 7.**
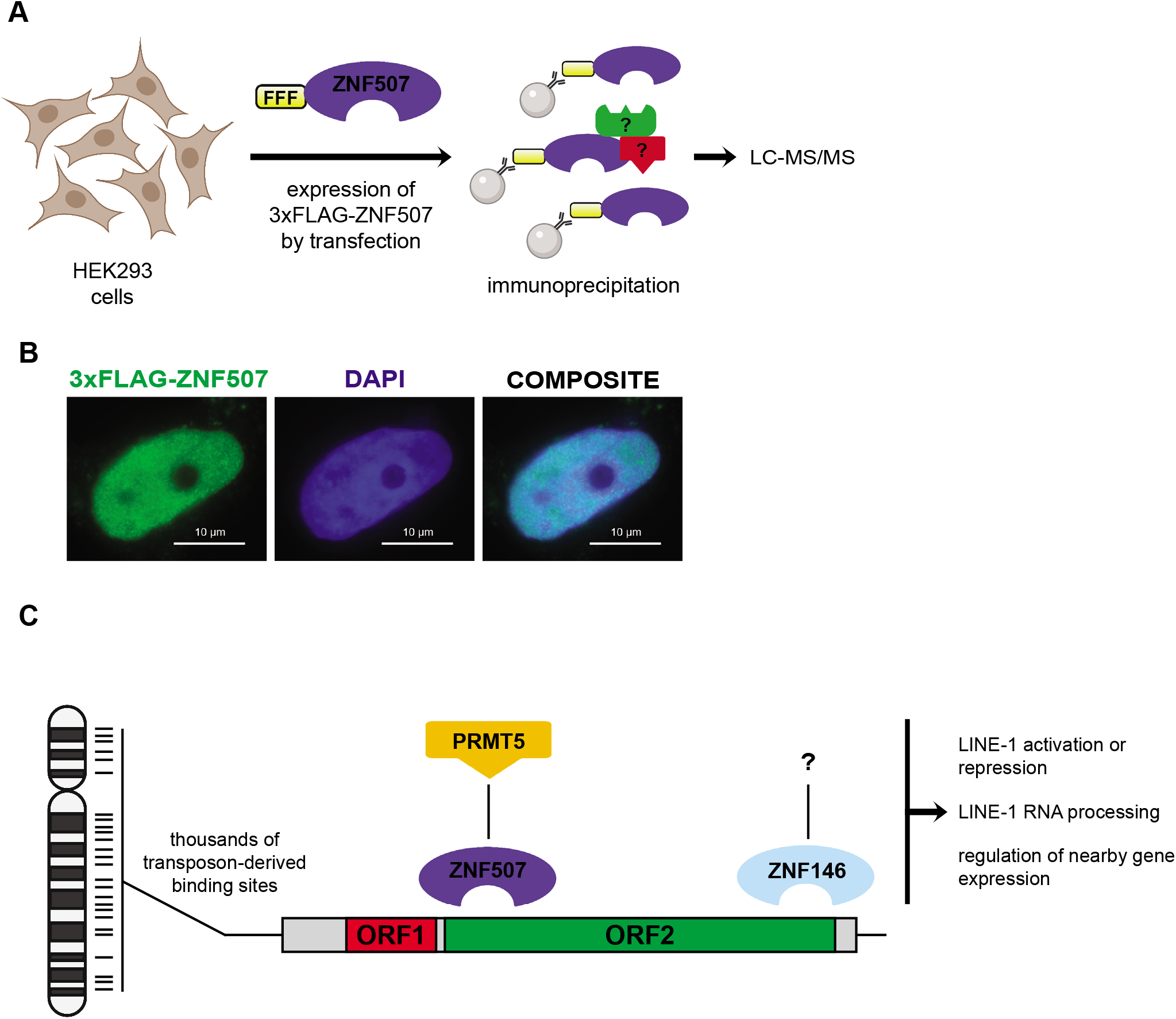
Identification of an interaction between ZNF507 and PMRT5. (A) Schematic of approach to identify ZNF507-interacting proteins. HEK293 cells were transiently transfected with an 3xFLAG tagged ZNF507 expression plasmid or an empty vector control. After 48 hours proteins were extracted and subjected to immunoprecipitation using anti-FLAG beads. Proteins were eluted by competition with 3xFLAG peptide before limited short gel electrophoresis and analysis by mass spectrometry (LC-MS/MS). (B) Immunofluorescence analysis of HEK293 cells expressing 3xFLAG-ZNF507. All interphase cells observed expressing 3xFLAG-ZNF507 had nuclear localization. No appreciable signal was observed for cells transfected with empty vector 3xFLAG control plasmid. (C) Model for LINE-1 binding by ZNF146 and ZNF507 and potential functions. ZNF146 and ZNF507 bind at thousands of sites in the genome as a result of LINE-1 transposition. Binding by either protein may then recruit various activities to these regions, such as the potential for ZNF507 to recruit protein arginine methylation activity through interaction with PRMT5. These activities may then contribute to context or cell-type dependent regulation of LINE-1 elements or nearby genes. Schematics created with BioRender.com.

As expected, bait protein, ZNF507, had the highest peptide count in the 3xFLAG-ZNF507 IP with far more peptides in this sample compared to empty vector control (Table 2). The second most abundant protein (by peptide count) was protein arginine methyltransferase 5 (PRMT5), again with far more peptides detected in the 3xFLAG-ZNF507 IP than for the EV control. Interestingly, PRMT5 is a known transcriptional and post-transcriptional regulator of gene expression with various protein targets including core histones (Bedford and Clarke, 2009; Dacwag et al., 2007; Lacroix et al., 2008; LeBlanc et al., 2016; Tee et al., 2010). These results suggest ZNF507 physically associates with PRMT5 and may recruit arginine methylation to L1 sequences to regulate L1 or endogenous gene expression (see model, Figure 7C).

**Table 2.** Results of co-IP LC-MS/MS.

| **Protein** | **# peptides\*** |  |
| --- | --- | --- |
|  | **3xFLAG-ZNF507** | **EV** |
| ZNF507 | 285 | 11 |
| PRMT5 | 45 | 12 |
| VCP | 36 | 5 |
| PPM1B | 35 | 11 |
| JUP | 33 | 14 |
| PRKDC | 33 | 0 |
| ANKFY1 | 28 | 5 |
| DSG1 | 23 | 10 |
| EEF1A1 | 22 | 5 |
| SLC25A5 | 22 | 1 |
| STK38 | 21 | 5 |
| EIF4B | 20 | 4 |
| RBM10 | 20 | 5 |
| AMY1A | 19 | 8 |
| EEF1G | 19 | 3 |
| ROCK1 | 19 | 1 |
| POF1B | 19 | 1 |
| LMNA | 18 | 0 |
| ACAP2 | 17 | 1 |
| WDR77 | 16 | 13 |
| ANXA1 | 16 | 12 |
| ANXA2 | 16 | 7 |
| EIF3B | 16 | 7 |
| KCTD17 | 16 | 5 |
| ENO1 | 16 | 1 |

\*Results represent the most abundant proteins (by number of peptides observed) after LC-MS/MS analysis of immunoprecipitations from HEK293 cells transfected with either 3xFLAG-ZNF507 or EV control expression plasmids as indicated. Common contaminants: keratins, heat shock proteins, tubulins, DSPN, FLNA, TBB5, T complex were excluded from this table.

## Discussion

About half of the human genome is comprised of repeat sequences that may serve underexplored roles in coordinated genome regulation. Unearthing proteins that target repetitive sequences may therefore serve as a launching pad for greater understanding of such processes. Over the past decade thousands of next generation data sets have been produced and made publicly available, most of which were processed without repeat sequences in mind. Taking a repeat-centric approach to mining these data sets may facilitate better understanding of the repeat genome and its influence on the biology of higher organisms.

There are many potential approaches to studying repeat enrichment in next generation sequencing data sets (Teissandier et al., 2019). Here we took a simple peak intersection approach and analyzed publish ENCODE data sets to identify proteins specifically enriched on repetitive transposon-derived sequences. We identified over 200 TF ChIP-seq peak sets enriched on transposon sequences more than would be expected by random chance. We then focused on a prominent enrichment of two putative L1-targeting proteins, ZNF146/OZF and ZNF507, and characterized their relationship to L1 sequences.

Other groups have previously taken similar approaches to identify proteins enriched on repetitive elements in ChIP-seq data sets, either using related peak intersection approaches or taking additional steps to extract information from reads mapping to multiple places in the genome (Barazandeh et al., 2018; Criscione et al., 2014; Jacobs et al., 2014; Schmitges et al., 2016; Soraya Shehata, 2020). Schmitges et al. used a similar approach to that taken here to calculate the percentage of peaks for ENCODE ChIP-seq data sets with transposable elements and independently identified ZNF146 as being enriched on L1 elements. During the preparation of this manuscript, ZNF507 was also recently identified by Shehata et al. as binding at L1 elements while performing large-scale analysis of ChIP-seq data sets.

Here we focus on ZNF146 and ZNF507 targeting to L1 sequences and find these proteins target different regions of the ORF2-encoding region of L1 DNA sequences in the human genome. ZNF146 targets both “old” and “young” L1 near the extreme 3’ end of ORF2. In contrast, ZNF507 was found to bind the 5’ end of L1 ORF2 and was only observed at younger elements. Thus, our findings add to the knowledge base of proteins that specifically recognize L1 elements in the genome including other L1-targeting proteins shown to differentiate between L1 subfamilies (Jacobs et al., 2014). We show that changes in the amino acid composition of ZNF507 zinc fingers during primate evolution paralleled changes to L1 DNA sequence, indicating ZNF507 and L1 sequence may have co-evolved during this time period or perhaps that ZNF507 may have recently acquired affinity for L1 repeats.

Surprisingly little is known about the functions of either ZNF146/OZF or ZNF507 despite being widely expressed in human tissue (Uhlen et al., 2015; Uhlen et al., 2017). Several years ago ZNF146 was shown to be overexpressed in various cancers, though the contribution or molecular mechanisms related to ZNF146/OZF in oncogenic transformation remains opaque (Antoine et al., 2005b; Ferbus et al., 1999; Ferbus et al., 2003; Ma et al., 2020; Zhu et al., 2021). ZNF146 was shown to associate with telomeric protein Rap1 and thus may have a role in telomere maintenance (Antoine et al., 2005a). Recently, ZNF146 was additionally shown to associate with replisome subunits and influence replication fork progression under conditions of DNA replication stress (Feu et al., 2020). Future studies are needed to determine how LINE-1 targeting and these potential roles in regulating genomic integrity at telomeres and during replication are related.

Although we did not observe broad upregulation of L1 elements or identify any gene expression changes in HEK293 cells lacking ZNF146, it remains possible that ZNF146 has important roles for regulating transcription or epigenetic modifications in heterochromatic regions, which are enriched for L1 sequences. If so, their contribution to regulation may be obscured by DNA methylation or other redundant silencing mechanisms, analogous to the maintenance of inactive-X heterochromatin despite removal of XIST RNA (Csankovszki et al., 1999). Similarly, ZNF146 could have a more essential role in specific developmental states, as it was recently identified as a potential regulator of the naïve primate pluripotency network (Boroviak et al., 2018). Since LINE-1 elements are known to be activated and regulate chromatin dynamics during early embryogenesis (Fadloun et al., 2013; Jachowicz et al., 2017), it’s tempting to speculate ZNF146/OZF is either involved in this activation or important for subsequent repression of LINE-1 elements post-activation. Future studies may address these potential connections between ZNF146, LINE-1 elements, pluripotency, and early development.

ZNF507 has been identified in genomic studies as being a potential risk locus for neural developmental disorders (Curtis and Consortium, 2016; Talkowski et al., 2012). Despite this potential relationship, to our knowledge ZNF507 had not been the subject of directed investigation. Our experiments here demonstrate ZNF507 targets L1 “young” elements and indicate ZNF507 depletion influences gene expression, potentially through interaction with PRMT5. Our transcriptomic analysis was performed in an embryonic kidney cell line (HEK293), which may not be the most relevant cell type to study ZNF507 function. Future experiments depleting ZNF507 in relevant neural cell types or neural organoids are needed to determine whether ZNF507 acts as a transcriptional regulator in these contexts. Identifying these targets could be key in determining what impact, if any, ZNF507 has on various brain disorders.

PRMT5 was recently shown to be a critical repressor of LINE-1 elements in primordial germ cells and early embryonic development (Kim et al., 2014). In this context, PRMT5-mediated methylation of histones is required to repress L1 expression during this unusual period of global reduction in DNA methylation. To our knowledge, how PRMT5 is targeted to LINE-1 sequences is largely unknown. Here we identify a potential interaction between PRMT5 and ZNF507. Although this relationship requires further validation, results here indicate ZNF507’s primary function may be to recruit PRMT5-mediated arginine methyltransferase activity to chromatin in order to regulate L1 or endogenous gene expression during development.

---

**Supplemental Table 1.** Enrichment analysis of ENCODE ChIP-seq peak sets. Columns contain the ratio of observed peak intersection versus expected (average of randomly shuffled peaks) for each TE class.

## Materials and Methods

### TE Enrichment Analysis

All TF ChIP-seq experiments with a corresponding IDR threshold peak file were downloaded from ENCODE in March 2020 (hg19 only). For each experiment, peaks were shuffled 10 times using BEDtools shuffle excluding ENCODE blacklist regions and -noOverlapping -maxTries 1000 parameters (Quinlan and Hall, 2010). Frequency of intersection between experimental or shuffled peaks and Repeatmasker repeat classes were calculated using BEDtools intersect and the -u parameter. Heatmaps were graphed in R.

### LINE-1 enrichment analysis

Meta-analysis of ChIP enrichment at LINE-1 elements was performed using unique reads (fold change over control bigwig files downloaded from ENCODE). Primary alignments were also downloaded from ENCODE and coverage files of fold change over control coverage were also generated without filtering for uniqueness using deepTools bamCoverage and the – normalizeTo1X parameters (Ramirez et al., 2014). Meta-analysis was performed at full-length L1 elements annotated by L1Base or binned by subfamily from Repeatmasker (Penzkofer et al., 2017). Mappability tracks were downloaded from the UCSC table browser. Heatmaps were generated using deepTools computeMatrix and plotHeatmap software.

Coverage of peaks aligned to the L1PA1 consensus was performed by extracting reads (ENCODE) mapping to L1 elements using BEDtools intersect. Aligned sequences were then retrieved using BEDtools getfasta. Fasta sequences were then directly aligned to the L1PA1 consensus sequence using bwa mem and the -B 1 -O 1 -d 1 -T 1 -r 0.1 -t 12 -k 10 parameters (Li and Durbin, 2009). Coverage was then calculated using Samtools depth. Signal over input was then calculated and graphed in R using ggplot2.

### Motif discovery and scanning

Motif discovery was performed on ENCODE peak DNA sequences using MEME software and order-0 background, classic discovery mode, zero or one occurrence, 13mer parameters (Bailey and Elkan, 1994). Motif occurrences in the genome were found using the top identified motif for each ChIP experiment and FIMO software, filtered by the indicated p-value cutoff (Grant et al., 2011).

### Zinc finger identification and multiple sequence alignments

Protein sequences were downloaded for alignment from UniProt. L1 sequences were downloaded from Dfam. Zinc fingers were predicted and annotated using online software (http://zf.princeton.edu/) (Persikov et al., 2009). Multiple sequence alignments were performed using ClustalX and Jalview with additional manual curation to focus on zinc finger residues and presentation.

### CRISPR/Cas9 disruption and RNA sequencing

HEK293 cells were grown in DMEM (Thermo Fisher) media and transiently transfected with equal amounts of Cas9 and sgRNA expressing plasmids (PX459) targeting either ZNF146, ZNF507 or an equivalent amount of empty vector control plasmid (Ran et al., 2013). Target sequences used for ZNF146 were 5’-ACTGAGCATGAGCATTTTC-3’, 5’-ACATGTACAATAAGTGATG-3’, 5’-ACTGTAAATTCTCTGCTGGC-3’, and 5’-TGAAGGTTTTTCCACACTC-3’. Target sequences used for ZNF507 were 5’-TGGGCTTCAAGTTCCTCC-3’, 5’-ATTGTTTCCGGACAAACTT-3’, 5’-CTGGCTTCTAGATGTAATA-3’, and 5’-AGGTTGGCTCTTGTCAACTC-3’. The day after transfection cells were selected for 36 hours using puromycin before outgrowth for a total of two weeks prior to analysis.

Western blots were performed using antibodies against ZNF146 (Novus Biologicals) or ZNF507 (Thermo Fisher). RNA was isolated from cells disrupted with CRISPR/Cas9 in biological duplicate using Trizol RNA extraction reagent. RNA was treated with TURBO DNase (Thermo Fisher) to remove contaminating DNA and cleaned up using RNeasy Mini (Qiagen) kits according to manufacturer recommendations. RNA sequencing libraries were prepared using a stand specific kit RNA-seq kit with ribosome depletion (KAPA Biosystems) and NEBNext Adapters and Multiplex Oligos for Illumina. Paired-end 50 bp sequencing was performed on the Illumina HiSeq 2000 platform (UMMS Core Facilities).

Reads were pre-processed and clipped using the Fastx Toolkit and those mapping to ribosomal RNA were removed prior to subsequent mapping with Bowtie. Non-ribosomal reads were then mapped to the human genome (hg19) using TopHat2 software with --library-type fr-firststrand -no-coverage-search parameters and Ensembl (ver82) gene model annotations. Differential expression analysis was performed using count matrices generated by featureCounts and DeSeq2 (Liao et al., 2014). Associated graphs were generated using ggplot2 within R. The associated fastq files and count matrices for this experiment are available at GEO: GSE172285.

### Expression, immunofluorescence analysis, and immunoprecipitation of FLAG-tagged ZNF507

To express 3xFLAG-tagged ZNF507, RNA ZNF507 cDNA was amplified from HeLa cell RNA using NEBNext^®^ High-Fidelity 2X PCR Master Mix and the following oligos: 5’-GATCGATCGGATCCGAAGAAAGTAGCAGTGTTGCCATGTTGGTG-3’ and 5’-GATCGATCGAATTCCTAATTTGTGTTTAGAGCTGTATTGTGGTCCTTATTCAGG-3’. ZNF507 cDNA was then and cloned into a pcDNA3.1 n-terminal 3xFLAG expression vector (CMV promoter).

For immunofluorescence analysis, HEK293 cells were grown on coverslips in 6 well dishes and transfected with pcDNA3.1-3XFLAG-ZNF507 plasmid using lipofectamine reagent (Thermo Fisher). After 36 hours cells were fixed with 4% paraformaldehyde in PBS for 10 minutes at room temperature. Coverslips were then incubated with a 1:100 dilution of M2 anti-FLAG antibody (sigma) in 1% BSA, 1X PBS at 37°C for 1 hour, then washed, and immunodetected using 1:500 dilution of conjugated secondary antibody, in 1X PBS with 1% BSA. Nuclei were visualized using Axiovert 200 microscope equipped with a 100X PlanApo objective (NA 1.4) and Chroma 83000 multibandpass dichroic and emission filter set (Bratteboro). Images were captured using a cooled charge-coupled device (CCD) camera (200 series, Photometrics). Images were minimally corrected for brightness and contrast using standard practices to best represent signals observed by eye using Zen (Zeiss) software. In cells with signal, 3xFLAG-ZNF507 consistently was localized to the nucleus. No appreciable signal was observed for cells transfected with empty vector control.

Immunoprecipitation of 3xFLAG-ZNF507 and associated proteins was performed by transfecting approximately 40 million HEK293 cells with either 3x-FLAG-ZNF507 expression plasmid or an empty vector control. After 36 hours, cells were dislodged from flasks in ice cold 1X PBS and harvested in 10 ml falcon tubes. Cells were washed twice in ice cold PBS before lysis for 5 minutes on ice in 1ml Lysis Buffer (10mM HEPES pH 7.5, 1.5mM MgCl2, 10mM NaCl, 0.075% NP-40, 1mM PMSF, 1X EDTA-free protease inhibitor cocktail (Roche)). Crude nuclear pellets were then harvested by centrifugation at 4°C for 3 min at 1000 x g and removal of supernatant. Nuclei were then suspended in 1ml Nuclear Extraction Buffer (20mM Tris-HCl pH7.5, 500mM NaCl, 1.5 mM MgCl2, 25% glycerol, 1mM PMSF, 1X EDTA-free protease inhibitor cocktail (Roche)), vortexed briefly, and incubated 15 minutes on ice. Insoluble DNA and debris were then cleared from the nuclear extract by centrifugation at 4°C for 10 minutes at 12,000 x g. To the supernatant an equal amount of Dilution buffer (20mM Tris-HCl pH 7.5, 1mM EDTA, 0.2% Triton-X 100, 1mM PMSF, 1X EDTA-free protease inhibitor cocktail (Roche)) was added to lower salt concentration before immunoprecipitation.

Nuclear extracts were pre-cleared by incubation with 40 μl Protein A dynabeads (Sigma) for one hour at 4°C. After capture of beads, 40 μl anti-FLAG M2 magnetic beads (Sigma) were added to each nuclear extract and incubated 6 hours at 4°C. Beads and co-immunoprecipitated proteins were then captured and washed twice for 5 minutes with rotation at 4°C with Wash buffer (20mM Tris-HCl pH 7.5, 150mM NaCl) containing 0.1% Triton X-100 and once without detergent. Proteins were eluted twice using 50 μl of 3X-FLAG peptide at a concentration of 500 ng/ μl and incubated with rotation at 4°C for 30 minutes for each elution.

Laemmli sample buffer (4X) was added to each elution to 1X final concentration and samples were incubated for 15 minutes at 80°C. Samples were then briefly run on the same 4–20% Criterion Stain Free™ Tris-HCl Protein Gel separated by several lanes so that the dye front was 2 cm below each well. Lanes were cut out slightly above the dye to the well bottom for LC-MS/MS analysis (UMMS Mass Spectrometry Core Facility). Mascot was set up to search SwissProt_Human and Scaffold (Proteome Software Inc.) was used to validate MS/MS based peptide and protein identifications.

## Acknowledgements

The authors would like to acknowledge other members of the Lawrence lab for feedback and support with the preparation of this manuscript. This work was supported by the NIH R35 GM122597.

## Author Contributions

This study was conceptualized by K.M.C. and J.B.L. Experiments and analysis were performed by K.M.C. and E.C.L. The manuscript was written by K.M.C. with input from E.C.L. under the supervision of J.B.L.

## References

Antoine, K., Ferbus, D., Kolahgar, G., Prosperi, M.T., and Goubin, G. (2005a). Zinc finger protein overexpressed in colon carcinoma interacts with the telomeric protein hRap1. J Cell Biochem 95, 763–768.

Antoine, K., Prosperi, M.T., Ferbus, D., Boule, C., and Goubin, G. (2005b). A Kruppel zinc finger of ZNF 146 interacts with the SUMO-1 conjugating enzyme UBC9 and is sumoylated in vivo. Mol Cell Biochem 271, 215–223.

Bailey, T.L., and Elkan, C. (1994). Fitting a mixture model by expectation maximization to discover motifs in biopolymers. Proc Int Conf Intell Syst Mol Biol 2, 28–36.

Barazandeh, M., Lambert, S.A., Albu, M., and Hughes, T.R. (2018). Comparison of ChIP-Seq Data and a Reference Motif Set for Human KRAB C2H2 Zinc Finger Proteins. G3 (Bethesda) 8, 219–229.

Bedford, M.T., and Clarke, S.G. (2009). Protein arginine methylation in mammals: who, what, and why. Mol Cell 33, 1–13.

Boroviak, T., Stirparo, G.G., Dietmann, S., Hernando-Herraez, I., Mohammed, H., Reik, W., Smith, A., Sasaki, E., Nichols, J., and Bertone, P. (2018). Single cell transcriptome analysis of human, marmoset and mouse embryos reveals common and divergent features of preimplantation development. Development 145.

Bourque, G., Leong, B., Vega, V.B., Chen, X., Lee, Y.L., Srinivasan, K.G., Chew, J.L., Ruan, Y., Wei, C.L., Ng, H.H., et al. (2008). Evolution of the mammalian transcription factor binding repertoire via transposable elements. Genome Res 18, 1752–1762.

Brouha, B., Schustak, J., Badge, R.M., Lutz-Prigge, S., Farley, A.H., Moran, J.V., and Kazazian, H.H., Jr. (2003). Hot L1s account for the bulk of retrotransposition in the human population. Proc Natl Acad Sci U S A 100, 5280–5285.

Burns, K.H. (2017). Transposable elements in cancer. Nat Rev Cancer 17, 415–424.

Consortium, E.P. (2012). An integrated encyclopedia of DNA elements in the human genome. Nature 489, 57–74.

Cosby, R.L., Chang, N.C., and Feschotte, C. (2019). Host-transposon interactions: conflict, cooperation, and cooption. Genes Dev 33, 1098–1116.

Coufal, N.G., Garcia-Perez, J.L., Peng, G.E., Yeo, G.W., Mu, Y., Lovci, M.T., Morell, M., O’Shea, K.S., Moran, J.V., and Gage, F.H. (2009). L1 retrotransposition in human neural progenitor cells. Nature 460, 1127–1131.

Criscione, S.W., Zhang, Y., Thompson, W., Sedivy, J.M., and Neretti, N. (2014). Transcriptional landscape of repetitive elements in normal and cancer human cells. BMC Genomics 15, 583.

Csankovszki, G., Panning, B., Bates, B., Pehrson, J.R., and Jaenisch, R. (1999). Conditional deletion of Xist disrupts histone macroH2A localization but not maintenance of X inactivation. Nat Genet 22, 323–324.

Curtis, D., and Consortium, U.K. (2016). Practical Experience of the Application of a Weighted Burden Test to Whole Exome Sequence Data for Obesity and Schizophrenia. Ann Hum Genet 80, 38–49.

Dacwag, C.S., Ohkawa, Y., Pal, S., Sif, S., and Imbalzano, A.N. (2007). The protein arginine methyltransferase Prmt5 is required for myogenesis because it facilitates ATP-dependent chromatin remodeling. Mol Cell Biol 27, 384–394.

Deininger, P., Morales, M.E., White, T.B., Baddoo, M., Hedges, D.J., Servant, G., Srivastav, S., Smither, M.E., Concha, M., DeHaro, D.L., et al. (2017). A comprehensive approach to expression of L1 loci. Nucleic Acids Res 45, e31.

Dewannieux, M., Esnault, C., and Heidmann, T. (2003). LINE-mediated retrotransposition of marked Alu sequences. Nat Genet 35, 41–48.

Ewing, A.D., and Kazazian, H.H., Jr. (2010). High-throughput sequencing reveals extensive variation in human-specific L1 content in individual human genomes. Genome Res 20, 1262–1270.

Fadloun, A., Le Gras, S., Jost, B., Ziegler-Birling, C., Takahashi, H., Gorab, E., Carninci, P., and Torres-Padilla, M.E. (2013). Chromatin signatures and retrotransposon profiling in mouse embryos reveal regulation of LINE-1 by RNA. Nat Struct Mol Biol 20, 332–338.

Faulkner, G.J., and Garcia-Perez, J.L. (2017). L1 Mosaicism in Mammals: Extent, Effects, and Evolution. Trends Genet 33, 802–816.

Ferbus, D., Antoine, K., and Goubin, G. (1999). Production and characterization of mouse monoclonal antibodies to human zinc finger OZF protein overexpressed in pancreatic carcinomas. Hybridoma 18, 431–436.

Ferbus, D., Bovin, C., Validire, P., and Goubin, G. (2003). The zinc finger protein OZF (ZNF146) is overexpressed in colorectal cancer. J Pathol 200, 177–182.

Feu, S., Unzueta, F., Llopis, A., Semple, J.I., Ercilla, A., Guaita-Esteruelas, S., Jaumot, M., Freire, R., and Agell, N. (2020). OZF is a Claspin-interacting protein essential to maintain the replication fork progression rate under replication stress. FASEB J 34, 6907–6919.

Giordano, J., Ge, Y., Gelfand, Y., Abrusan, G., Benson, G., and Warburton, P.E. (2007). Evolutionary history of mammalian transposons determined by genome-wide defragmentation. PLoS Comput Biol 3, e137.

Grant, C.E., Bailey, T.L., and Noble, W.S. (2011). FIMO: scanning for occurrences of a given motif. Bioinformatics 27, 1017–1018.

Hall, L.L., Carone, D.M., Gomez, A.V., Kolpa, H.J., Byron, M., Mehta, N., Fackelmayer, F.O., and Lawrence, J.B. (2014). Stable C0T-1 repeat RNA is abundant and is associated with euchromatic interphase chromosomes. Cell 156, 907–919.

Imbeault, M., Helleboid, P.Y., and Trono, D. (2017). KRAB zinc-finger proteins contribute to the evolution of gene regulatory networks. Nature 543, 550–554.

Jachowicz, J.W., Bing, X., Pontabry, J., Boskovic, A., Rando, O.J., and Torres-Padilla, M.E. (2017). LINE-1 activation after fertilization regulates global chromatin accessibility in the early mouse embryo. Nat Genet 49, 1502–1510.

Jacobs, F.M., Greenberg, D., Nguyen, N., Haeussler, M., Ewing, A.D., Katzman, S., Paten, B., Salama, S.R., and Haussler, D. (2014). An evolutionary arms race between KRAB zinc-finger genes ZNF91/93 and SVA/L1 retrotransposons. Nature 516, 242–245.

Kazazian, H.H., Jr., and Moran, J.V. (2017). Mobile DNA in Health and Disease. N Engl J Med 377, 361-370.

Khan, H., Smit, A., and Boissinot, S. (2006). Molecular evolution and tempo of amplification of human LINE-1 retrotransposons since the origin of primates. Genome Res 16, 78–87.

Kigami, D., Minami, N., Takayama, H., and Imai, H. (2003). MuERV-L is one of the earliest transcribed genes in mouse one-cell embryos. Biol Reprod 68, 651–654.

Kim, S., Gunesdogan, U., Zylicz, J.J., Hackett, J.A., Cougot, D., Bao, S., Lee, C., Dietmann, S., Allen, G.E., Sengupta, R., et al. (2014). PRMT5 protects genomic integrity during global DNA demethylation in primordial germ cells and preimplantation embryos. Mol Cell 56, 564–579.

Kolosha, V.O., and Martin, S.L. (1997). In vitro properties of the first ORF protein from mouse LINE-1 support its role in ribonucleoprotein particle formation during retrotransposition. Proc Natl Acad Sci U S A 94, 10155–10160.

Konkel, M.K., Walker, J.A., and Batzer, M.A. (2010). LINEs and SINEs of primate evolution. Evol Anthropol 19, 236–249.

Korenberg, J.R., and Rykowski, M.C. (1988). Human genome organization: Alu, lines, and the molecular structure of metaphase chromosome bands. Cell 53, 391–400.

Lacroix, M., El Messaoudi, S., Rodier, G., Le Cam, A., Sardet, C., and Fabbrizio, E. (2008). The histone-binding protein COPR5 is required for nuclear functions of the protein arginine methyltransferase PRMT5. EMBO Rep 9, 452–458.

Laity, J.H., Lee, B.M., and Wright, P.E. (2001). Zinc finger proteins: new insights into structural and functional diversity. Curr Opin Struct Biol 11, 39–46.

LeBlanc, S.E., Wu, Q., Lamba, P., Sif, S., and Imbalzano, A.N. (2016). Promoter-enhancer looping at the PPARgamma2 locus during adipogenic differentiation requires the Prmt5 methyltransferase. Nucleic Acids Res 44, 5133–5147.

Li, H., and Durbin, R. (2009). Fast and accurate short read alignment with Burrows-Wheeler transform. Bioinformatics 25, 1754–1760.

Liao, Y., Smyth, G.K., and Shi, W. (2014). featureCounts: an efficient general purpose program for assigning sequence reads to genomic features. Bioinformatics 30, 923–930.

Lu, J.Y., Chang, L., Li, T., Wang, T., Yin, Y., Zhan, G., Han, X., Zhang, K., Tao, Y., Percharde, M., et al. (2021). Homotypic clustering of L1 and B1/Alu repeats compartmentalizes the 3D genome. Cell Res.

Ma, Y., Cong, X., Zhang, Y., Yin, X., Zhu, Z., and Xue, Y. (2020). CircPIP5K1A facilitates gastric cancer progression via miR-376c-3p/ZNF146 axis. Cancer Cell Int 20, 81.

Manuelidis, L., and Ward, D.C. (1984). Chromosomal and nuclear distribution of the HindIII 1.9-kb human DNA repeat segment. Chromosoma 91, 28–38.

Mathias, S.L., Scott, A.F., Kazazian, H.H., Jr., Boeke, J.D., and Gabriel, A. (1991). Reverse transcriptase encoded by a human transposable element. Science 254, 1808–1810.

Mir, A.A., Philippe, C., and Cristofari, G. (2015). euL1db: the European database of L1HS retrotransposon insertions in humans. Nucleic Acids Res 43, D43–47.

Moran, J.V., Holmes, S.E., Naas, T.P., DeBerardinis, R.J., Boeke, J.D., and Kazazian, H.H., Jr. (1996). High frequency retrotransposition in cultured mammalian cells. Cell 87, 917–927.

Ostertag, E.M., Goodier, J.L., Zhang, Y., and Kazazian, H.H., Jr. (2003). SVA elements are nonautonomous retrotransposons that cause disease in humans. Am J Hum Genet 73, 1444–1451.

Penzkofer, T., Jager, M., Figlerowicz, M., Badge, R., Mundlos, S., Robinson, P.N., and Zemojtel, T. (2017). L1Base 2: more retrotransposition-active LINE-1s, more mammalian genomes. Nucleic Acids Res 45, D68–D73.

Persikov, A.V., Osada, R., and Singh, M. (2009). Predicting DNA recognition by Cys2His2 zinc finger proteins. Bioinformatics 25, 22–29.

Philippe, C., Vargas-Landin, D.B., Doucet, A.J., van Essen, D., Vera-Otarola, J., Kuciak, M., Corbin, A., Nigumann, P., and Cristofari, G. (2016). Activation of individual L1 retrotransposon instances is restricted to cell-type dependent permissive loci. Elife 5.

Quinlan, A.R., and Hall, I.M. (2010). BEDTools: a flexible suite of utilities for comparing genomic features. Bioinformatics 26, 841–842.

Ramirez, F., Dundar, F., Diehl, S., Gruning, B.A., and Manke, T. (2014). deepTools: a flexible platform for exploring deep-sequencing data. Nucleic Acids Res 42, W187–191.

Ran, F.A., Hsu, P.D., Wright, J., Agarwala, V., Scott, D.A., and Zhang, F. (2013). Genome engineering using the CRISPR-Cas9 system. Nat Protoc 8, 2281–2308.

Rebollo, R., Romanish, M.T., and Mager, D.L. (2012). Transposable elements: an abundant and natural source of regulatory sequences for host genes. Annu Rev Genet 46, 21–42.

Robbez-Masson, L., and Rowe, H.M. (2015). Retrotransposons shape species-specific embryonic stem cell gene expression. Retrovirology 12, 45.

Schmitges, F.W., Radovani, E., Najafabadi, H.S., Barazandeh, M., Campitelli, L.F., Yin, Y., Jolma, A., Zhong, G., Guo, H., Kanagalingam, T., et al. (2016). Multiparameter functional diversity of human C2H2 zinc finger proteins. Genome Res 26, 1742–1752.

Smit, A.F., Toth, G., Riggs, A.D., and Jurka, J. (1995). Ancestral, mammalian-wide subfamilies of LINE-1 repetitive sequences. J Mol Biol 246, 401–417.

Soraya Shehata, S.S., Alison Swearingen, Graycen Wheeler, Arpan Das, Giulia Corbet, Benjamin Nebenfuehr, Daniel Ahrens, Devin Tauber, Shelby Lennon, Kevin Choi, Thao Huynh, Tom Weiser, Kristen Schneider, Michael Bradshaw, Maria Lai, Joel Basken, Tim Read, Jon Demasi, Matt Hynes-Grace, Dan Timmons, Michael Smallegan, John L. Rinn (2020). Genome-wide binding analysis of 195 DNA Binding Proteins reveals “reservoir” promoters and human specific SVA-repeat family regulation. Biorxiv.

Talkowski, M.E., Rosenfeld, J.A., Blumenthal, I., Pillalamarri, V., Chiang, C., Heilbut, A., Ernst, C., Hanscom, C., Rossin, E., Lindgren, A.M., et al. (2012). Sequencing chromosomal abnormalities reveals neurodevelopmental loci that confer risk across diagnostic boundaries. Cell 149, 525–537.

Tee, W.W., Pardo, M., Theunissen, T.W., Yu, L., Choudhary, J.S., Hajkova, P., and Surani, M.A. (2010). Prmt5 is essential for early mouse development and acts in the cytoplasm to maintain ES cell pluripotency. Genes Dev 24, 2772–2777.

Teissandier, A., Servant, N., Barillot, E., and Bourc’his, D. (2019). Tools and best practices for retrotransposon analysis using high-throughput sequencing data. Mob DNA 10, 52.

Uhlen, M., Fagerberg, L., Hallstrom, B.M., Lindskog, C., Oksvold, P., Mardinoglu, A., Sivertsson, A., Kampf, C., Sjostedt, E., Asplund, A., et al. (2015). Proteomics. Tissue-based map of the human proteome. Science 347, 1260419.

Uhlen, M., Zhang, C., Lee, S., Sjostedt, E., Fagerberg, L., Bidkhori, G., Benfeitas, R., Arif, M., Liu, Z., Edfors, F., et al. (2017). A pathology atlas of the human cancer transcriptome. Science 357.

Wang, H., Xing, J., Grover, D., Hedges, D.J., Han, K., Walker, J.A., and Batzer, M.A. (2005). SVA elements: a hominid-specific retroposon family. J Mol Biol 354, 994–1007.

Wang, J., Xie, G., Singh, M., Ghanbarian, A.T., Rasko, T., Szvetnik, A., Cai, H., Besser, D., Prigione, A., Fuchs, N.V., et al. (2014). Primate-specific endogenous retrovirus-driven transcription defines naive-like stem cells. Nature 516, 405–409.

Zhu, S., Chen, C.Y., and Hao, Y. (2021). LncRNA KCNQ1OT1 acts as miR-216b-5p sponge to promote colorectal cancer progression via up-regulating ZNF146. J Mol Histol.

